# SIP2 functions as the master transcription factor of the *Plasmodium* merozoite formation

**DOI:** 10.1101/2024.08.02.606280

**Authors:** Tsubasa Nishi, Izumi Kaneko, Masao Yuda

## Abstract

Malaria, one of the most serious infectious diseases worldwide, is caused by the proliferation of *Plasmodium* parasites through repeated cycles of intraerythrocytic development. The parasite replicates via schizogony in host erythrocytes, producing multiple progeny merozoites that invade new erythrocytes to continue the intraerythrocytic developmental cycle. Although merozoite formation is the most crucial step in parasite proliferation and malaria pathogenesis, the molecular mechanism regulating merozoite formation remains unclear. SIP2 is an AP2 transcription factor expressed during schizogony and is particularly conserved among erythrocyte-infecting apicomplexan parasites. Here, we reveal that SIP2 in *P. berghei* (PbSIP2) functions as a transcriptional activator that regulates merozoite formation. Disruption of *pbsip2* using a dimerizable Cre recombinase system resulted in developmental arrest before merozoite formation and significant downregulation of merozoite-related genes. ChIP-seq of PbSIP2 showed that it comprehensively activated merozoite-related genes by binding to previously reported *cis*-regulatory elements of merozoite invasion-related genes, including the bipartite motif (TGCAN_4-6_GTGCA). Collectively, our results indicate that SIP2 is a transcription factor that establishes erythrocyte infectivity and may have an evolutionary origin from the common ancestor of erythrocyte-infecting apicomplexan parasites.

## Introduction

*Plasmodium* parasites replicate asexually in the blood of vertebrate hosts, which causes malaria, one of the most serious diseases worldwide, with more than 200 million cases and 600 thousand deaths every year^1^. The proliferation of parasites begins with a merozoite invading the host red blood cells (RBCs). Within RBC, parasites grow and begin a unique mode of cell division called schizogony^2^, wherein the parasites develop into multinucleated cells, or schizonts, through several asynchronous rounds of DNA replication and nuclear division (karyokinesis)^3–5^. Eventually, multiple merozoites are produced through segmentation (cytokinesis) simultaneously with the final round of karyokinesis^6^. The progeny merozoites then egress from the RBC and invade a new RBC to continue with IDC. Schizogony, the final step in an intraerythrocytic developmental cycle (IDC), is a crucial process for parasite proliferation in the host blood and is thus strongly related to parasite pathogenesis. However, the molecular mechanisms that regulate schizogony is still largely unknown.

The transcriptional regulation during the *Plasmodium* IDC has been assessed in several time-course transcriptomic studies^7–10^. These studies indicated periodic expression of the majority of genes during IDC. Incidentally, the time-course assay for transposase-accessible chromatin (ATAC)-seq further showed the transition of active promoter regions along with periodic transcriptomes^11^. For the schizont stage in particular, those transcriptomic studies demonstrated stage-specific expression of genes essential for major merozoite-specific structures, such as rhoptry and pellicle/inner membrane complex (IMC). Furthermore, ATAC-seq demonstrated the enrichment of several DNA motifs within regulatory regions that are open during schizogony. These motifs include a putative *cis*-regulatory element upstream of the invasion-related genes, GGTGCA (named PfM18.1), which was detected through an *in silico* search of DNA motifs in *P. falciparum*^12^. In addition, the *in silico* study identified a bipartite motif comprising GTGCA and GTGCA-like motifs upstream of genes encoding rhoptry proteins and suggested conservation of this bipartite motif in *Plasmodium* spp. These studies indicate the existence of a stage-specific transcription factor that regulates merozoite-related genes at the schizont stage by recognizing these motifs. However, such transcription factors have not yet been identified, and the molecular mechanisms regulating the transcriptome during schizogony remain unclear.

AP2 transcription factors are the major family of transcription factors found in *Plasmodium*^13^. Members in this family contain sequence-specific DNA binding domains with three anti-parallel β-sheets and one α-helix, called AP2 domain^14^. SIP2 is an AP2 transcription factor expressed during the schizont stage. In *P. falciparum*, SIP2 (PfSIP2; PF3D7_0604100) has been reported to bind to the subtelomeric var promoter element 2 (SPE2)^15^ and has been proposed to be involved in chromosome end biology^16^. However, the primary functions of SIP2 in *Plasmodium* remain to be elucidated. In the present study, we investigated the role of SIP2 in *P. berghei* (PbSIP2; PBANKA_0102900) during schizont development. We revealed that PbSIP2 plays an essential role in the final step of schizogony, *i.e.* merozoite formation, by activating the majority of genes important for merozoite-specific structures and abilities. Our data further showed that PbSIP2 was associated with previously reported *cis*-regulatory motifs of invasion-related genes, including the bipartite motif^12^.

## Results

### PbSIP2 is an AP2-family transcription factor that contains tandem AP2 domains conserved in RBC-infecting parasites

PbSIP2 contains two tandem AP2 domains at its N-terminus and a putative nuclear localization signal (NLS) on the C-terminal side of the AP2 domains (Fig 1A). Alignment of amino acid (AA) sequences among AP2 domains of PbSIP2 and its orthologs in other *Plasmodium* species showed that the two AP2 domains were conserved by 88 and 69%, respectively (Fig 1B). In addition, the AA sequences of the linker between the two AP2 domains were also conserved, suggesting that these two AP2 domains could cooperatively bind to genomic DNA (Fig 1B).

**Fig 1.**
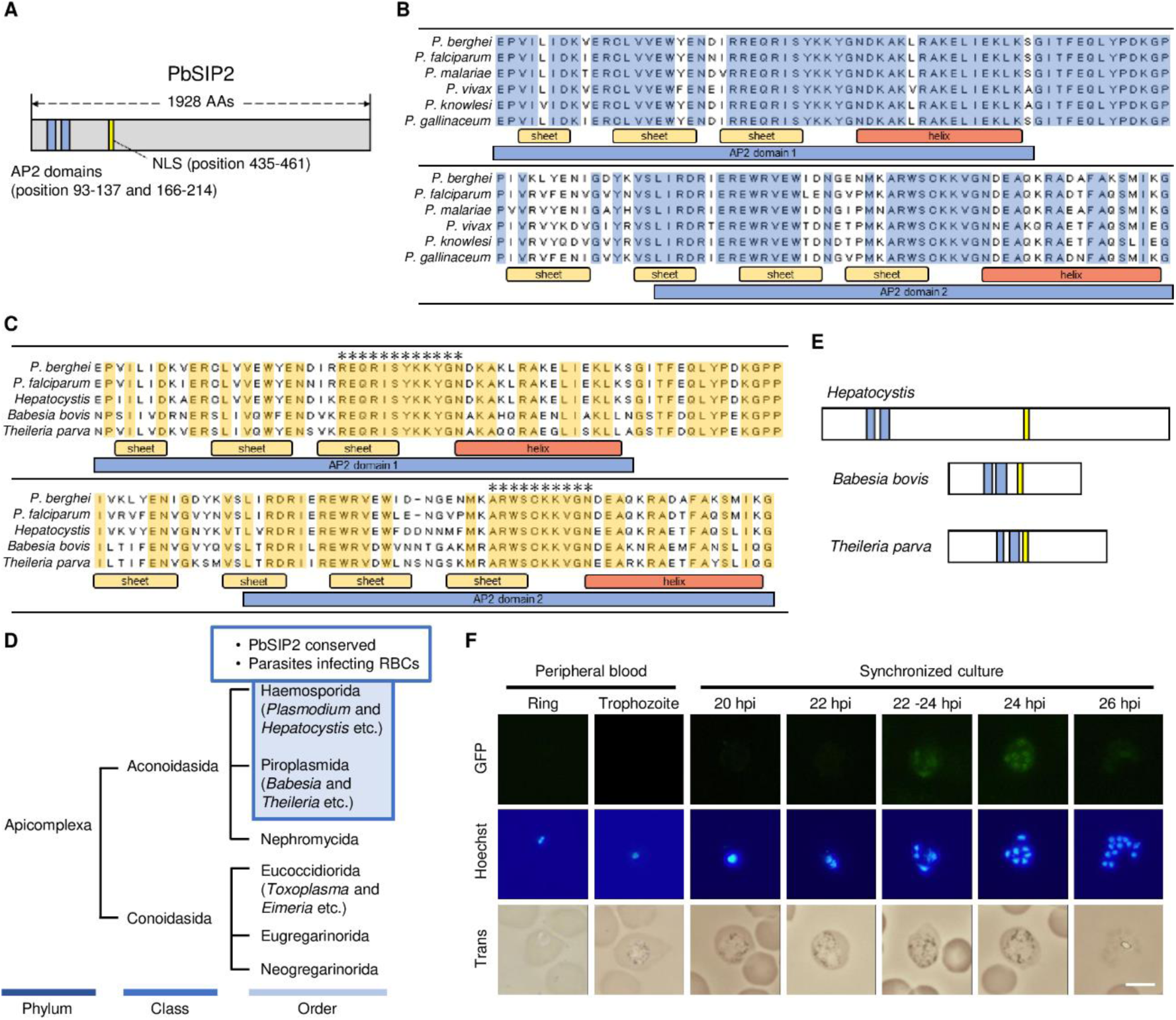
Sequence features of PbSIP2 and its expression during asexual-blood stage development. (A) Schematic illustration of PbSIP2. Blue boxes show the AP2 domains. The nuclear localization signal (NLS) was predicted using cNLS Mapper (http://nls-mapper.iab.keio.ac.jp/cgi-bin/NLS_Mapper_form.cgi) and indicated using a yellow box. (B) Alignment of amino acid sequences of PbSIP2 orthologs in *Plasmodium* by the ClustalW program in Mega X (*P*. *berghei*, PbANKA_0102900; *P*. *falciparum*, PF3D7_0604100; *P. malariae*, PmUG01_11058700; *P. vivax*, PVP01_1144800; *P*. *knowlesi*, PKNH_1146400; *P. gallinaceum*, PGAL8A_00132600). Amino acid sequences conserved in all aligned sequences are indicated by blue boxes. Yellow and red bars indicate a beta-sheet and alpha-helix, respectively. Blue bars show a sequence for an AP2 domain. (C) Alignment of amino acid sequences of PbSIP2, PfSIP2, and BLASTP-detected proteins (*Hepatocystis* spp., VWU52144; *Babesia Bovis*, XP_001610230; *Theileria Parva*, XP_764727). Amino acid sequences conserved in all aligned sequences are indicated by yellow boxes. Regions wherein more than 10 amino acids are continuously conserved are indicated with asterisks. (D) Taxonomic tree of Apicomplexa. Orders in which PbSIP2 is conserved are indicated in a blue box. (E) Schematic illustration of putative SIP2 orthologs in Apicomplexa. Blue boxes show the AP2 domains, and NLSs are indicated using a yellow box. (F) Expression of PbSIP2 in asexual blood-stage of the PbSIP2::GFP. Rings and trophozoites were detected from the peripheral blood, and schizonts were observed in synchronized cultures. Nuclei were stained with Hoechst 33342. Scale bar = 5 μm.

To further explore the conservation of PbSIP2 in *Apicomplexa*, we performed a protein-protein BLAST (BLASTP) search using the AA sequence of its AP2 domains and the linker between them (positions 87–214) as a query. The search detected proteins from *Hepatocystis*, *Babesia*, and *Theileria* species with E-values < 10^-50^, but not from other apicomplexan species (Fig 1C). Notably, *Plasmodium* and these PbSIP2-conserved species belong to the Haemosporida or Piroplasmida, which constitute a group of parasites that infect host RBCs^17,18^ (Fig 1D). AA sequences of the N- and C-terminal side AP2 domains were conserved with 59 and 62%, respectively, and regions between the third β-sheet and α-helix were completely conserved for both AP2 domains (Fig 1C). In addition, the linker lengths between the two AP2 domains were the same (28 AAs) for PbSIP2 and the BLASTP-detected proteins (Fig 1C). Furthermore, their overall structures were also similar to those of PbSIP2, as they contained the AP2 domains near their N-terminus and a putative NLS towards their C-terminal side (Fig 1E).

### PbSIP2 is expressed during the early schizont development

To determine the stage at which PbSIP2 functions, we generated a parasite line expressing GFP-fused PbSIP2 (PbSIP2::GFP, Fig S1) and assessed its expression pattern during the asexual blood stage. In the fluorescence analysis of the peripheral blood, no fluorescent signals were observed in the ring or trophozoite stages of PbSIP2::GFP (Fig 1F). The analysis was further performed for the schizont stages in culture because schizonts do not appear in the peripheral blood owing to the sequestration of schizont-infected RBCs^19^. We synchronized the cell cycle of PbSIP2::GFP by intravenously injecting cultured mature schizonts into mice. The synchronized parasites were cultured again at 6 h post-injection (hpi), and fluorescent signals were assessed every 2 h. In the time-course analysis, fluorescent signals first appeared in the schizonts with four nuclei at 22 hpi (Fig 1F). The signals continued until the schizonts had eight nuclei and then faded in later stages (Fig 1F). *Plasmodium* schizonts undergo several rounds of asynchronous nuclear division and a final round of karyokinesis, which simultaneously occurs with segmentation (cytokinesis), to produce multiple merozoites. Moreover, in *P. berghei*, a median of 12–13 merozoites is formed per schizont, with a maximum of 20^20^. Thus, these results indicate that PbSIP2 functions from the early rounds of karyokinesis until the beginning of cytokinesis.

### Disruption of *pbsip2* affects the late stage of schizont development

To investigate the role of PbSIP2 during schizont development, we performed a conditional knockout of *pbsip2* using a dimerizable Cre recombinase (DiCre) system^21,22^. We first developed a parasite line constitutively expressing Cre59 (Thr19-Asn59) fused with FKBP12 and Cre60 (Asn60-Asp343) fused with FRB, using the Cas9 expressing parasite, PbCas9^23^ (PbDiCre, Fig S2A). Subsequently, two loxP sequences were inserted on 5’ and 3’ sides of *pbsip2*, arranged in the same direction (*pbsip2*-cKO, Fig 2A and S2B). This parasite will lose the entire open reading frame of *pbsip2* in the presence of rapamycin (Fig 2A).

**Fig 2.**
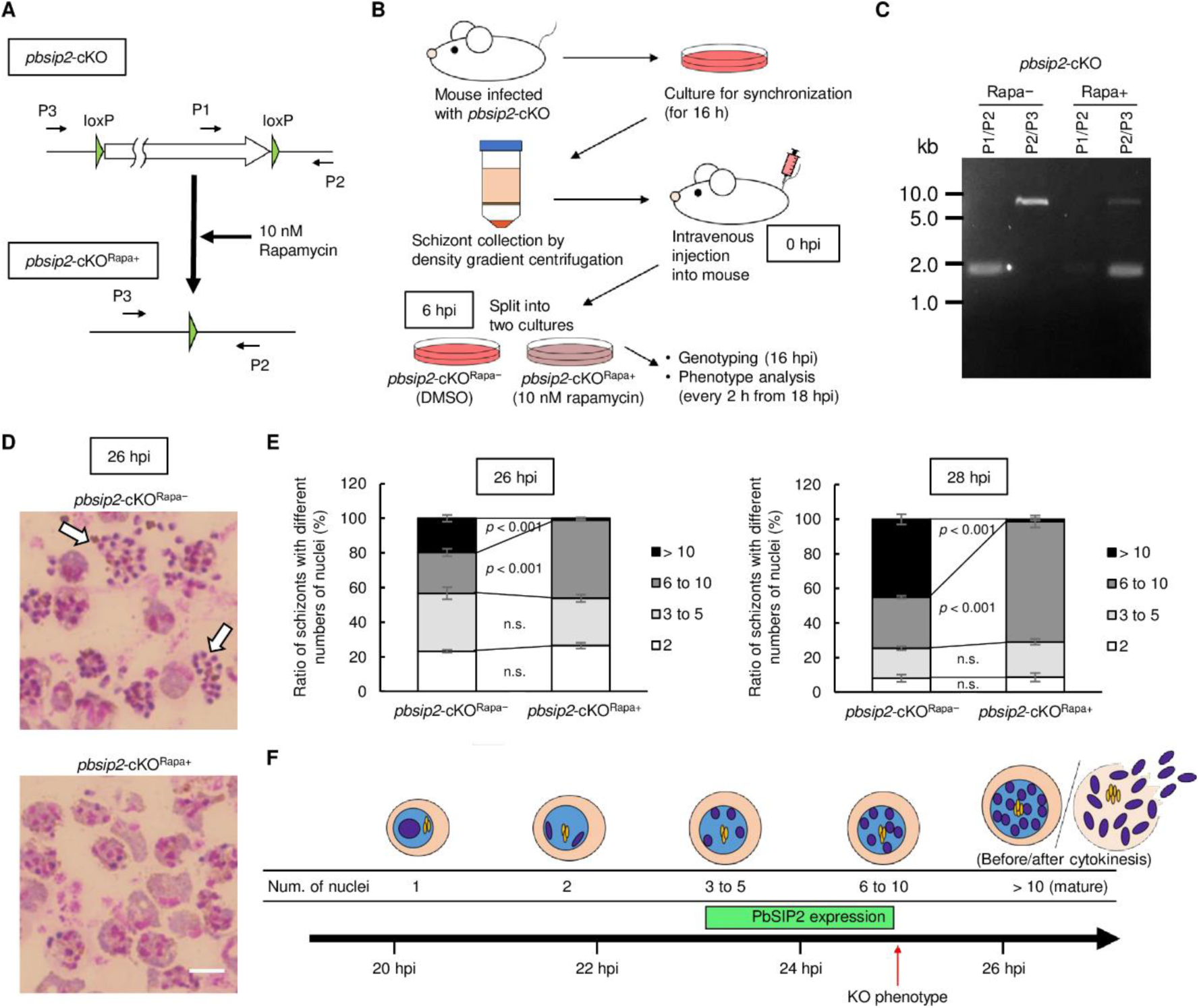
Conditional knockout of *pbsip2* by DiCre-mediated recombination. (A) Schematic illustration of DiCre-mediated recombination at the *pbsip2* locus of *pbsip2*-cKO. Recombination occurs between two loxP sites to excise *pbsip2* from the genome in the presence of rapamycin. P1–P3 indicate primers used for genotyping PCR. (B) Outline of conditional knockout experiments using *pbsip2*-cKO. First, the whole blood from mice infected with *pbsip2*-cKO was cultured. After 16 h of culture, schizonts were harvested by density gradient centrifugation and injected into mice. At 6 h post-injection (hpi), the whole blood was harvested and split into two cultures: one with 10 nM rapamycin and the other with DMSO. (C) A gel image of the genotyping PCR analysis for *pbsip2*-cKO^Rapa−^ and *pbsip2*-cKO^Rapa+^ performed at 16 hpi. Primers used are illustrated in (A). (D) Representative Giemsa-stained images of *pbsip2*-cKO^Rapa−^ and *pbsip2*-cKO^Rapa+^ at 26 hpi. Mature schizonts with merozoites inside are indicated by arrows. Scale bar = 5 μm. (E) Ratio of schizonts with different numbers of nuclei in *pbsip2*-cKO^Rapa−^ and *pbsip2*-cKO^Rapa+^ at 26 hpi (left) and 28 hpi (right). Error bars indicate the standard error of mean value from three independent experiments. The *p*-values were calculated by two-tailed Student’s t-test (n. s. stands for not significant, *p*-value > 0.01). (F) Model of the *P. berghei* schizont development in synchronized cultures. The period for PbSIP2 expression is indicated as a green box.

We first assessed whether DiCre-mediated recombination in *pbsip2*-cKO occurs quickly enough to disrupt *pbsip2* before PbSIP2 expression begins. The whole blood from mice infected with synchronized *pbsip2*-cKO was split into two cultures at 6 hpi, and the one was treated with 10 nM rapamycin (*pbsip2*-cKO^Rapa+^) and the other with dimethyl sulfoxide as a control (*pbsip2*-cKO^Rapa−^) (Fig 2B). At 18 hpi (12 h of culture), when no parasites had started nuclear division, recombination at the *pbsip2* locus was assessed by genotyping PCR. The assay showed that the *pbsip2* locus was almost completely excised from the genome in *pbsip2*-cKO^Rapa+^ while no recombination was detected in *pbsip2*-cKO^Rapa−^ (Fig 2C). Notably, using the parental PbDiCre parasite, we confirmed that schizont development was not affected by rapamycin (Fig S2C).

To assess the impact of *pbsip2*-disruption on schizont development, we observed *pbsip2*-cKO on Giemsa-stained smears every two hours from 18 hpi. In culture, *pbsip2*-cKO^Rapa+^ developed as comparable to *pbsip2*-cKO^Rapa−^ until 24 hpi. At 26 hpi, approximately 20 % of *pbsip2*-cKO^Rapa−^ became mature schizonts (number of nuclei > 10), and some have already formed merozoites (Fig 2D and 2E). In contrast, most schizonts of *pbsip2*-cKO^Rapa+^ had less than ten nuclei at 26 hpi (Fig 2D and 2E). Furthermore, while the number of schizonts with six–ten nuclei increased at 28 hpi, mature schizonts were barely produced in *pbsip2*-cKO^Rapa+^, and no merozoite formation was observed (Fig 2D and 2E). This result suggests that the development of *pbsip2*-cKO^Rapa+^ is arrested before cytokinesis that accompanies the final round of karyokinesis (Fig 2F). Together with expression analysis, these results indicate that PbSIP2 plays an essential role in schizont development.

### PbSIP2 binds to the upstream of genes related to merozoite formation

Next, we performed chromatin immunoprecipitation followed by high-throughput sequencing (ChIP-seq) analysis using PbSIP2::GFP and investigated its binding sites on a genome-wide basis. PbSIP2::GFP parasites were synchronized as described above, and ChIP-seq was performed at 24 hpi. Two biologically independent experiments were conducted, and their overall peak patterns were highly similar, suggesting high reproducibility of the ChIP-seq experiments (Fig 3A). Using macs2, 784 and 804 peaks were identified in Experiments 1 and 2, respectively, and 645 peaks (more than 80% of the peaks in Experiment 1) overlapped between them (Fig 3B and Tables S1A and S1B). To investigate the binding motif of PbSIP2, we assessed the enrichment of sequence motifs within 50 bp of the summits of the common peaks using Fischer’s exact test. The analysis showed enrichment of GTGCA with a *p*-value less than 5.0 × 10^−324^ (the smallest positive real number on the R platform, hereafter referred to as *p*-value^limit^) (Fig 3C). The GTGCA motif was found within 300 bp of the summit of 556 peaks (86% of the common peaks), and the closer the motif was to the peak summit, more peaks were detected (Fig 3D). These results indicate that the GTGCA motif is the binding motif of PbSIP2.

**Fig 3.**
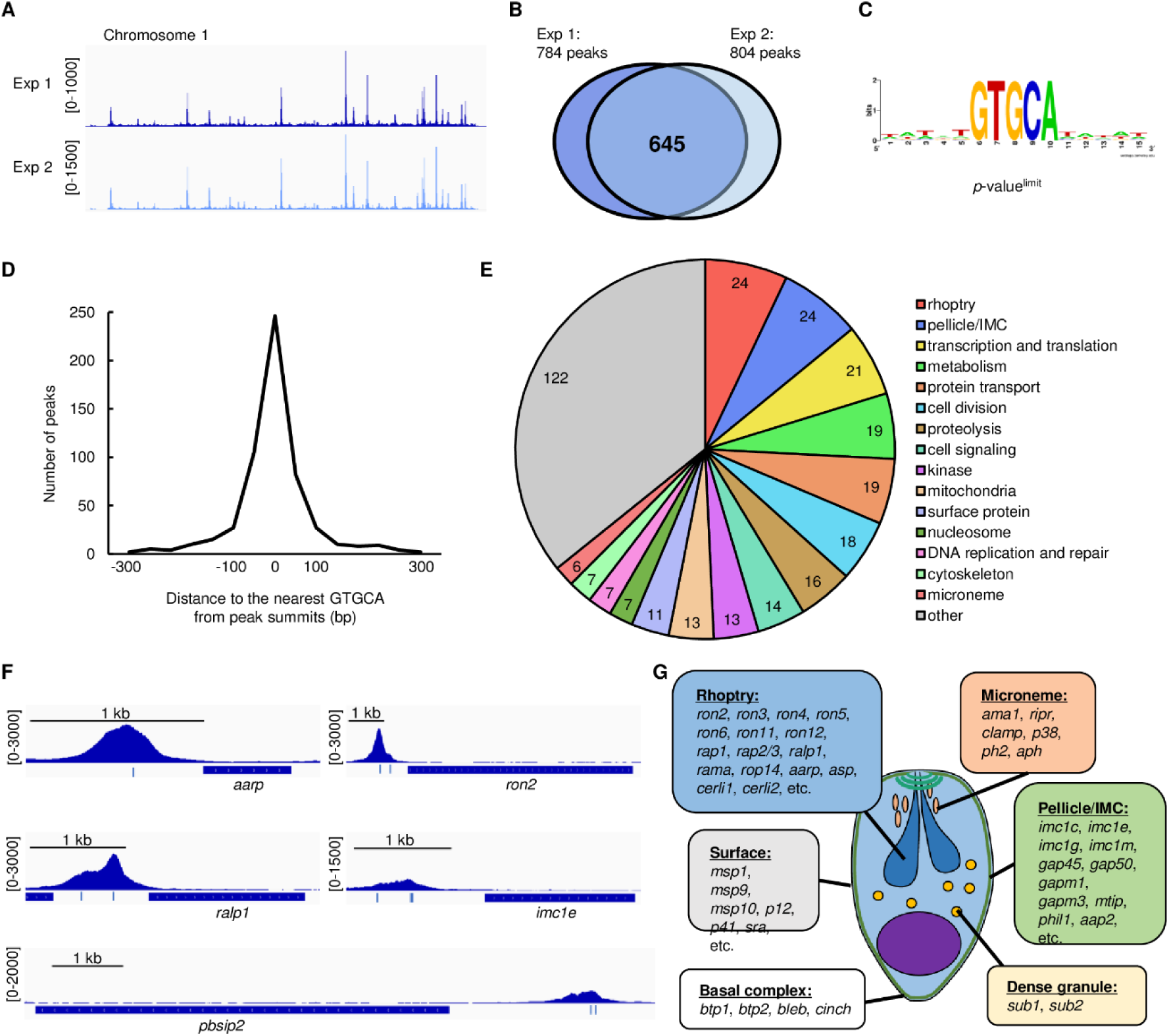
Chromatin immunoprecipitation followed by high-throughput sequencing (ChIP-seq) analysis of PbSIP2. (A) Integrative Genomics Viewer images for ChIP-seq Experiment 1 and 2 of PbSIP2 on the chromosome 1. Histograms show row read coverage of ChIP data at each base. Scales are indicated in square brackets. (B) A Venn diagram showing the number of overlapping peaks between Experiment 1 and 2. (C) A motif enriched within 50 bp from ChIP-seq peak summits of PbSIP2. The logo was depicted using WebLogo (https://weblogo.berkeley.edu/logo.cgi). *p*-value^limit^ is a *p*-value less than 5.0 × 10^−324^ (the smallest positive real number on the R platform). (D) Distance between peak summits identified in the ChIP-seq and the nearest GTGCA motifs. (E) Classification of functionally annotated target genes of PbSIP2 into 16 characteristic groups. The number of genes categorized into each group is indicated in the graph. (F) IGV images showing ChIP-seq peaks upstream of the PbSIP2 target genes. Histograms show row read coverage of ChIP data at each base. Locations of GTGCA motif are indicated as a blue bar. The scales are shown in square brackets. (G) Schematic illustration of *P. berghei* merozoite. Characteristic structures of the merozoite and PbSIP2 targets that belong to each structure are shown.

We next examined the PbSIP2 peak locations in the genome and identified target genes. By defining genes containing a peak within 1200 bp of their start codon as PbSIP2 targets, we identified 489 target genes from 645 common peaks (Table S1C). Of these, 340 are functionally annotated on PlasmoDB (https://plasmodb.org/plasmo/app). To evaluate the role of PbSIP2, we classified the annotated targets into 16 characteristic groups (Fig 3E). The analysis revealed that the two largest groups in the PbSIP2 targets were “rhoptry” and “pellicle/IMC.” The group “rhoptry” contained majority of known rhoptry genes, including the recently identified cytosolically exposed rhoptry leaflet interacting protein genes, *cerli1* and *cerli2*^24,25^ (Fig 3F and Fig 3G). The targets in “pellicle/IMC” included several glideosome-associated protein genes (such as *gap45* and *gap50*) and IMC protein genes (such as *imc1c* and *imc1e*) (Fig 3F and 3G). Moreover, the PbSIP2 targets contained genes related to other merozoite structures, such as microneme (such as *ama1* and *ripr*^26^), dense granules (such as *sub1* and *sub2*) merozoite surface (such as *msp1* and *msp9*), and basal complex (such as *btp1* and *bleb*^27^) (Fig 3G). These results demonstrated that PbSIP2 targets the majority of genes important for merozoite formation, which is consistent with the developmental arrest at the schizont stage observed in the conditional knockout of *pbsip2*. In addition, although not included among the PbSIP2 targets because of the upstream threshold of 1200 bp, *pbsip2* had a PbSIP2 peak approximately 2-kb upstream (Fig 3F). This result indicates that PbSIP2 could activate its gene through positive transcriptional feedback, which is characteristic of many transcriptional activators in *Plasmodium*^28–31^.

### PbSIP2 functions as a transcriptional activator

Next, we evaluated the role of PbSIP2 in transcriptional regulation during schizont development using high-throughput RNA sequencing (RNA-seq). Conditional knockout of *pbsip2* was performed as described above, and recombination was confirmed at 18 hpi. The *pbsip2*-cKO^Rapa+^ and *pbsip2*-cKO^Rapa−^ parasites were then subjected to RNA-seq analysis at 26 hpi, and the sequence data were compared between them using DESeq2 (Table S2). The analysis revealed that in *pbsip2*-cKO^Rapa+^, 193 genes were significantly downregulated [log_2_(fold change) < −1, *p*-value adjusted for multiple testing with the Benjamini-Hochberg procedure (*p*-value^adj^) < 0.05] compared to *pbsip2*-cKO^Rapa−^ [excluding *pbsip2*, which was downregulated with log_2_(fold change) of −4.0 and *p*-value^adj^ of 3.9 × 10^−231^] (Fig 4A). These significantly downregulated genes contained 115 target genes of PbSIP2, showing a significant overlap with a *p*-value of 7.9 × 10^−66^ by Fisher’s exact test (Fig 4B). In contrast, only two genes were significantly upregulated in *pbsip2*-cKO^Rapa+^ [log_2_(fold change) > 1, *p*-value^adj^ < 0.05], none of which were targets of PbSIP2 (Fig 4A). In addition, the two most enriched motifs in the upstream region (300– 1200 bp from ATG) of the downregulated genes contained the PbSIP2 binding motif (Fig 4C). These results indicated that PbSIP2 functions as a transcriptional activator during schizont development. The genes that were significantly downregulated in *pbsip2*-cKO^Rapa+^ included a considerable number of rhoptry genes (25 genes) (Fig 4D and 4E), 22 of which were also PbSIP2 target genes (Fig 4B). Furthermore, these rhoptry genes were tended to show higher −log_2_(fold change) values than the other downregulated genes; 8 of the 11 genes downregulated with log_2_(fold change) < −4 were a rhoptry gene (Fig 4E), suggesting that PbSIP2 functions as a strong *trans*-acting factor for transcriptional activation of rhoptry genes.

**Fig 4.**
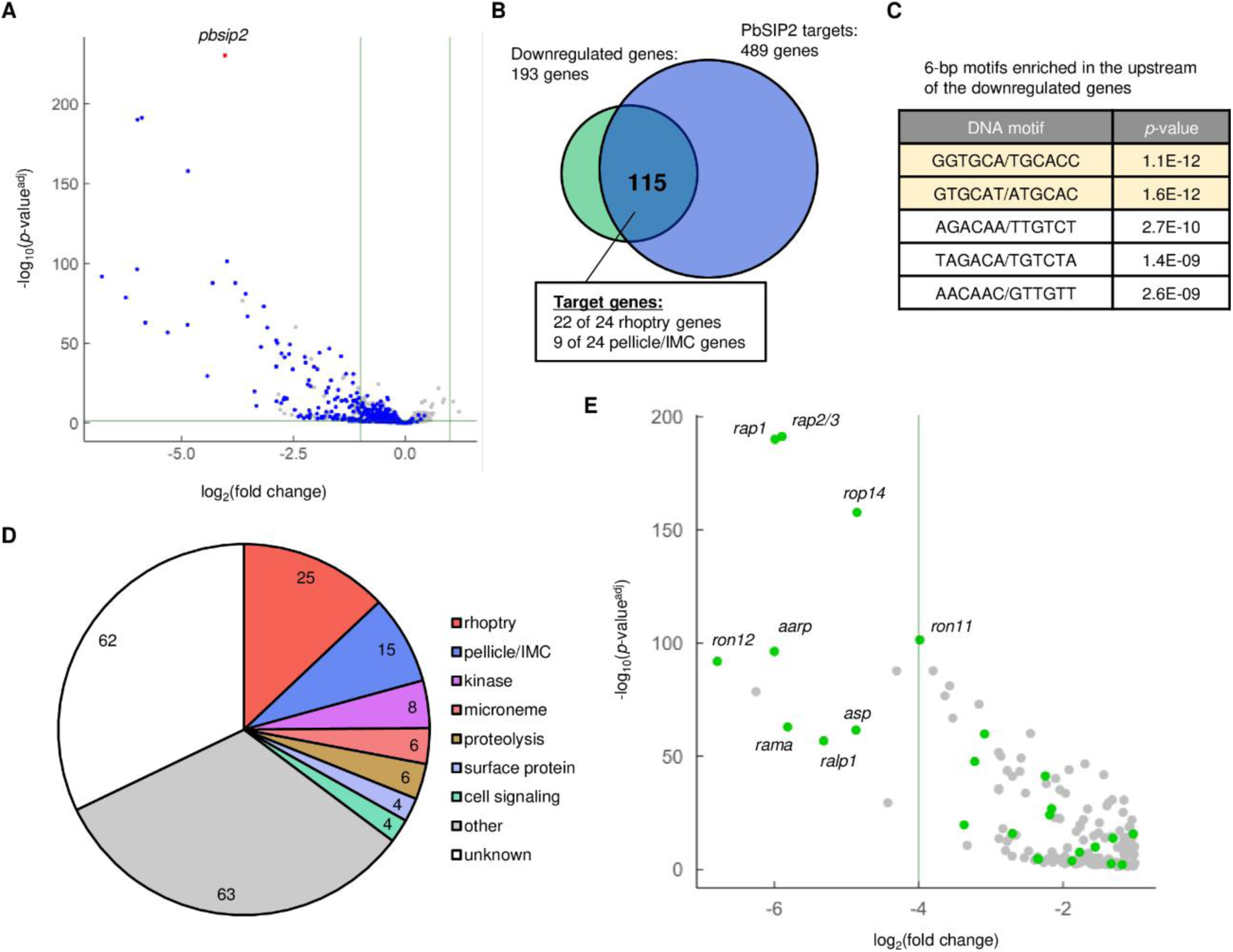
Differential expression analysis between *pbsip2*-cKO^Rapa−^ and *pbsip2*-cKO^Rapa+^ by RNA-seq. (A) Volcano plot showing differential expression of genes between *pbsip2*-cKO^Rapa−^ and *pbsip2*-cKO^Rapa+^. Blue dots represent target genes of PbSIP2, and a red dot represents *pbsip2*. A horizontal line indicates a *p*-value of 0.05 and two vertical lines indicate a log_2_(fold change) of 1 and −1. (B) A Venn diagram showing an overlap between the genes significantly downregulated in *pbsip2*-cKO^Rapa+^ and the PbSIP2 target genes. (C) Six-base motifs enriched in the upstream (300–1200 bp from start codons) of genes downregulated in *pbsip2*-cKO^Rapa+^. Motifs including GTGCA are indicated in light yellow. (D) Classification of the downregulated genes into characteristic groups. (E) A section of the Volcano plot from (A). Genes significantly downregulated in *pbsip2*-cKO^Rapa+^ compared with *pbsip2*-cKO^Rapa−^ are shown. Green dots represent a rhoptry gene, and names of some rhoptry genes are shown.

### PbSIP2 binds to the GTGCA motif through its tandem AP2 domains

To examine the DNA-binding properties of the AP2 domains of PbSIP2, we performed DNA immunoprecipitation followed by high-throughput sequencing (DIP-seq)^32^ using recombinant PbSIP2 AP2 domains fused with a maltose-binding protein tag on its N-terminal side (MBP::PbSIP2). MBP::PbSIP2 was mixed with fragmented *P. berghei* genomic DNA (approximately 150–300 bp). DNA fragments bound by MBP::PbSIP2 were harvested using amylose resin and sequenced by NGS. In the sequence data, we detected 3451 peaks throughout the genome (Fig 5A and Table S3). Within 50 bp from the summit of the DIP peaks, GTGCA was significantly enriched with a *p*-value^limit^ by Fischer’s exact test, similar to the ChIP-seq results. This result indicated that PbSIP2 directly binds to genomic DNA through its AP2 domains. In contrast to the ChIP-seq (Fig 3C), slight enrichment of G and A was observed on the 5’-side of GTGCA when sequence logo was constructed from the sequences around GTGCA detected in the DIP-seq peaks (Fig 5B). In fact, more than 70% of GTGCA motifs in the peak regions were RGTGCA (R = G or A). Consistently, in the motif enrichment analysis for 6-bp motifs, GGTGCA was the most enriched, followed by AGTGCA and motifs one-base-shifted from GGTGCA (Fig 5C). Therefore, the AP2 domains of PbSIP2 preferentially binds to RGTGCA. Notably, for the ChIP-seq peaks, those with RGTGCA showed a higher average fold enrichment value than the others with a *p*-value of 7.5 × 10^−6^ by two-tailed Student’s t-test (Fig 5D). Therefore, although enrichment of RGTGCA was not evident in ChIP-seq, PbSIP2 bound to RGTGCA with high affinity *in vivo*.

**Fig 5.**
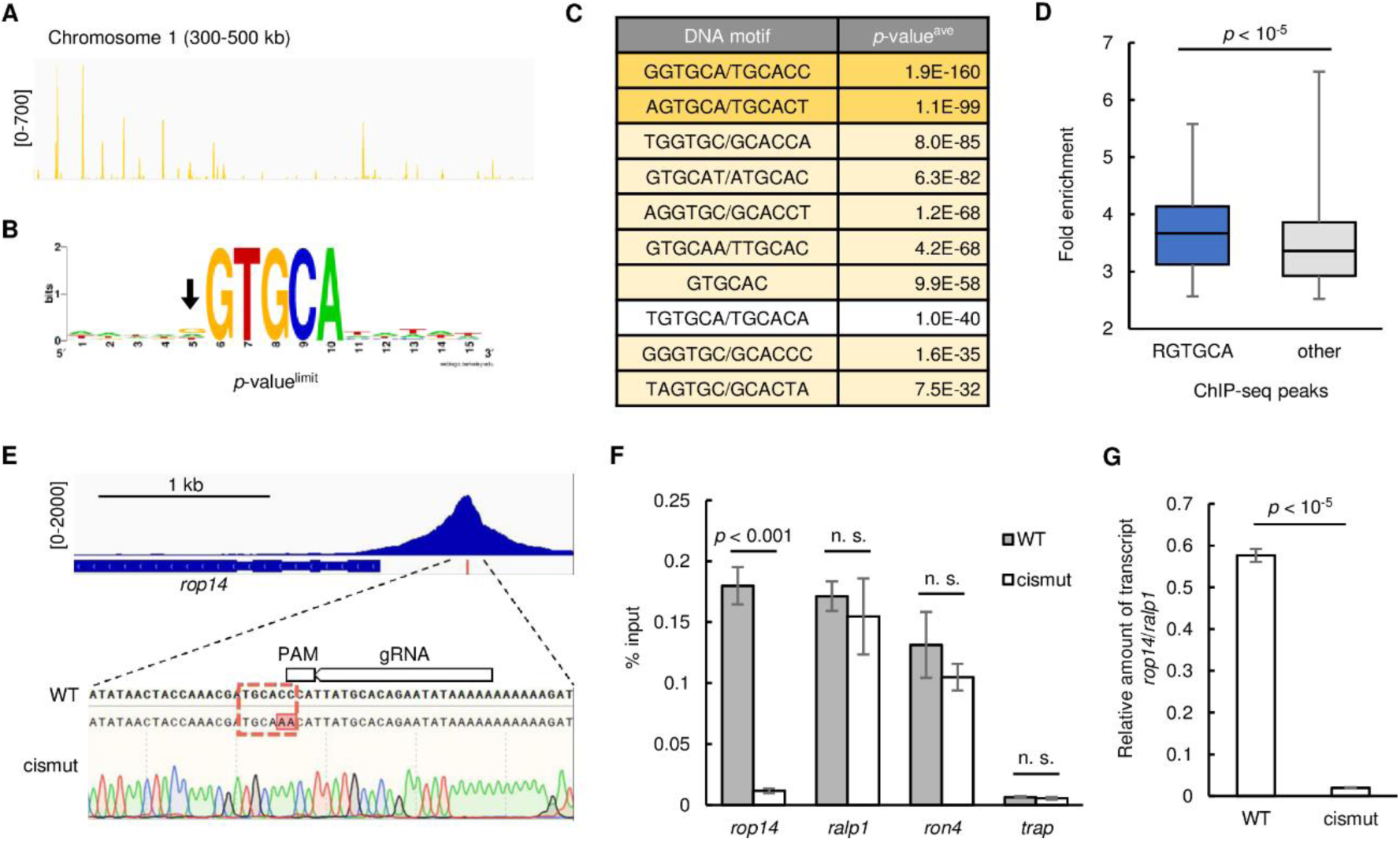
DNA-binding property of PbSIP2 and function of its binding motif as a *cis*-regulatory element. (A) An Integrative Genomics Viewer image for DNA immunoprecipitation followed by high-throughput sequencing (DIP-seq) using recombinant AP2 domains of PbSIP2 on the chromosome 1. Histograms show row read coverage of IP data at each base. Scales are indicated in square brackets. (B) A motif enriched within 50 bp from peak summits of DIP-seq. The logo was depicted using WebLogo. *p*-value^limit^ is a *p*-value less than 5.0 × 10^−324^ (the smallest positive real number on the R platform). An arrow indicates slight enrichment of G and A in the logo. (C) Six-base motifs enriched in the DIP-seq peaks. RGTGCA motif is indicated in yellow, and its one-base shifted motifs are indicated in light yellow. The enrichment analyses were separately performed for 7 chromosome sets (chromosome 1–4, 5–7, 8–9, 10–11, 12, 13, and 14) because the analysis using all DIP-seq peaks detected several motifs with *p*-value^limit^ (Fischer’s exact test yields a lower *p*-value when population becomes higher with a constant odds ratio). *p*-value^ave^ indicates the average *p*-values for each motif among the 7 analyses. Motifs were ranked along their *p*-value^ave^. (D) Box plot showing the distribution of fold enrichment values for the ChIP-seq peaks with RGTGCA (blue) and others (grey). The *p*-value was calculated by two-tailed Student’s t-test. (E) Integrative Genomics Viewer image showing the ChIP-seq peak in the upstream of *rop14* and genomic sequence around the RGTGCA motif in the peak region. Sanger sequence result for PbSIP2::GFP^Cas9_cismut^ is shown under the wild-type sequence. The target region of single guide RNA used for generating PbSIP2::GFP^Cas9_cismut^ is indicated on the sequence. (F) ChIP-qPCR analysis of PbSIP2 at the mutated site upstream of *rop14*. Error bars indicate the standard error of mean %input value from three independent experiments. The *p*-values were calculated by two-tailed Student’s t-test (n. s. stands for not significant, *p*-value > 0.01). [*rop14*: PBANKA_0111600, *ralp1*: PBANKA_0619700, *ron4*: PBANKA_0932000, *trap*: PBANKA_1349800] (G) RT-qPCR analysis of *rop14* transcripts between PbSIP2::GFP^Cas9^ and PbSIP2::GFP^Cas9_cismut^ parasites. Error bars indicate the standard error of mean value from three independent experiments. The *p*-value was calculated by two-tailed Student’s t-test.

### RGTGCA is essential for PbSIP2 binding and transcription of the downstream gene

To assess whether RGTGCA is essential for the DNA binding of PbSIP2 and functions as a *cis*-regulatory element for the downstream gene, we introduced a mutation into the motif within the ChIP peak upstream of *rop14* (Fig 5E). Binding of PbSIP2 and *rop14* transcription were assessed by ChIP coupled with quantitative PCR (ChIP-qPCR) and reverse transcription qPCR (RT-qPCR), respectively. ROP14 is a functionally uncharacterized rhoptry protein that localizes to the rhoptry bulbs of merozoites^33^. *rop14* was selected as the target in this experiment because it is not essential for the asexual blood-stage development in *P. berghei*. We developed parasites expressing GFP-fused PbSIP2 using PbCas9 (PbSIP2::GFP^Cas9^, Fig S3) and then introduced a mutation upstream of *rop14*, altering GGTGCA to ttTGCA (PbSIP2::GFP^Cas9_cismut^, Fig 5E). Using these parasite lines, we examined whether the disruption of the RGTGCA motif affected the binding of PbSIP2 to the genome by ChIP-qPCR analysis. In the PbSIP2::GFP^Cas9_cismut^ parasite, the amount of immunoprecipitated DNA fragments relative to the input DNA (%input value) upstream of *rop14* was more than 15-fold lower than that in PbSIP2::GFP^Cas9^ (Fig 5F). In contrast, upstream of other PbSIP2 targets, *ralp1* and *ron4*, the binding of PbSIP2 was not affected, as the %input values were comparable between the parasites with and without the mutation (Fig 5F). The %input value upstream of the non-target gene *trap* was assessed as a negative control and showed no significant change after the introduction of the mutation (Fig 5F).

Next, to investigate the function of the PbSIP2 binding motif as a *cis*-regulatory element, we compared the transcript levels of *rop14* between PbSIP2::GFP^Cas9^ and PbSIP2::GFP^Cas9_cismut^ by RT-qPCR analysis at 26 hpi. The transcriptional activity of the endogenous *rop14* promoter was assessed as the relative amount of *rop14* transcripts against *ralp1* transcripts. The analysis showed that in PbSIP2::GFP^Cas9_cismut^, the relative amount of *rop14* transcripts was approximately 30-fold lower than that in PbSIP2::GFP^Cas9^, revealing significant downregulation of *rop14* promoter activity after introducing the mutation (Fig 5G). These results demonstrate that RGTGCA is essential for PbSIP2 binding and functions as a *cis*-regulatory element for activating downstream genes.

### PbSIP2 recognize the bipartite motif upstream of rhoptry genes

A previous *in silico* study reported that two copies of the GTGCA (or GTGCA-like) motif, separated by five or six nucleotides, are conserved upstream of some rhoptry genes in *Plasmodium* species^12^. In the ChIP-seq of PbSIP2, GTGCA was identified as the binding motif, and most rhoptry genes were detected as targets. Considering these, we explored DNA sequences in the ChIP-seq peaks and detected bipartite motifs, in which GTGCA-like motifs were found on the 5’-side of GTGCA, upstream of some rhoptry genes (Fig 6A). Because PbSIP2 has two tandem AP2 domains, we hypothesized that PbSIP2 might recognize these bipartite motifs. Accordingly, we predicted the 3-D structure of the PbSIP2 AP2 domains using AlphaFold2 to evaluate whether they would have a structure capable of binding to two separate motifs^34^. In the predicted model, two AP2 domains were aligned in parallel, and the distance between them was 33.11 Å, which is close to the length per turn of B-DNA helix (34 Å) (Fig 6B). The 5-bp motifs separated by five or six nucleotides appear on the same side over one turn because the B-DNA helix is approximately 10 bp per turn (Fig 6B). Therefore, the two parallel AP2 domains seemed suitable for binding to the bipartite motifs upstream of rhoptry genes.

**Fig 6.**
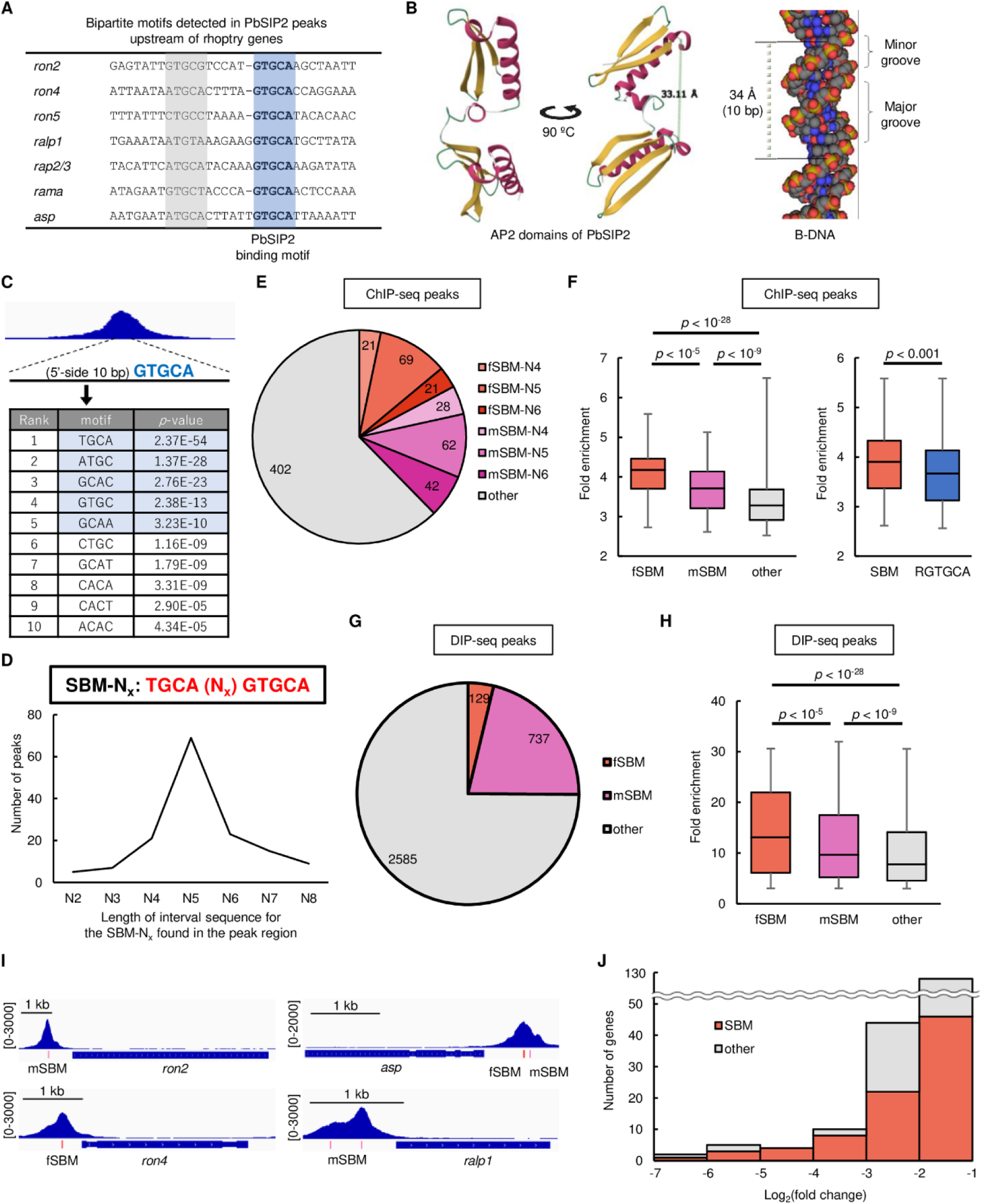
Association of PbSIP2 with schizont bipartite motif (SBM). (A) GTGCA-containing bipartite motifs detected in PbSIP2 peaks upstream of rhoptry genes. GTGCA and GTGCA-like motifs are indicated by blue and light green, respectively. (B) 3D structure of the PbSIP2 AP2 domains (position 87–214) predicted using AlphaFold2^34^. On the right, a B-DNA model depicted using a 3D molecule modeling tool GraphiteLifeExplorer^47^ is shown. (C) Four-bp motifs enriched within 5’-side 10 bp of GTGCA in the peak regions. The *p*-values were calculated by Fisher’s exact test. TGCA motif and its one-base-shifted motifs are indicated by blue. (D) The number of ChIP-seq peaks containing SBM-N_x_ with different length of interval sequence in the bipartite motif. (E) Pie graph showing the number of ChIP-seq peaks with fSBM (full SBM) and mSBM (SBM with a single mismatch in the 5’ TGCA). (F) Box plot showing distribution of fold enrichment values for the ChIP-seq peaks with fSBM (red), mSBM (pink), and others (grey). A box plot for the peaks with SBM (red) and RGTGCA (blue) is also shown on the right. The *p*-values were calculated by two-tailed Student’s t-test. (G) Pie graph showing the number of DIP-seq peaks containing fSBM and mSBM. (H) Box plot showing distribution of fold enrichment values for the DIP-seq peaks with fSBM (red), mSBM (pink), and others (grey). The *p*-values were calculated by two-tailed Student’s t-test. (I) Integrative Genomics Viewer images showing ChIP-seq peaks on the upstream of PbSIP2 target genes. Red and pink bars indicate location of the fSBM and mSBM, respectively. Scales are indicated in square brackets. (J) Histogram showing the number of genes against the log_2_(fold change) value in the differential expression analysis between *pbsip2*-cKO^Rapa−^ and *pbsip2*-cKO^Rapa+^. The number of genes that contain SBM in their upstream are indicated by red.

To further assess whether PbSIP2 is genome-widely associated with the bipartite motifs, we searched for enrichment of motifs within 10 bp on the 5’-side of GTGCA in the ChIP-seq peaks using Fischer’s exact test. The analysis detected TGCA as the most enriched motif with a *p*-value of 2.4 × 10^−54^ (Fig 6C). In addition, motifs one-base shifted from the TGCA, such as ATGC and GCAC, were detected with *p*-values < 1.0 × 10^−10^ (Fig 6C). Therefore, the results suggested that PbSIP2 could be associated with the bipartite motif having TGCA on the 5’-side of the GTGCA motif. Given this result, we next searched for peaks with TGCAN_x_GTGCA (schizont bipartite motif, SBM-N_x_) and detected 140 peaks containing SBM-N_2-8_ (Fig 6D). Among these peaks, those containing SBM-N_4-6_ were the majority (111 peaks), with the SBM-N_5_ peaks being the most abundant (Fig 6D). In addition, we also performed the search allowing a single mismatch in the 5’ TGCA and detected additional 132 peaks having the SBM-N_4-6_ with one mismatch (Fig 6E). Thus, nearly 40% of the PbSIP2 peaks are associated with these bipartite motifs. Hereafter, SBM-N_4-6_ and those containing a single mismatch are called fSBM (full SBM) and mSBM, respectively, and altogether called SBM.

The peaks with the SBM tended to have higher intensity compared with the other peaks; *i.e.*, average fold enrichment values of peaks with fSBM and mSBM were significantly higher than that of the other peaks with *p*-values of 2.7 × 10^−29^ and 5.4 × 10^−10^ by two-tailed Student’s t-test, respectively (Fig 6F). In addition, the average fold enrichment value of peaks with fSBM was higher than that of peaks with mSBM (*p*-value = 8.7 × 10^−6^), suggesting that the DNA-binding of PbSIP2 is stronger without any mismatches in the 5’ TGCA (Fig 6F). Notably, the peaks with SBM also had a higher average fold enrichment value than those with RGTGCA (*p*-value = 2.6 × 10^−4^) (Fig 6F). Collectively, these results indicate that PbSIP2 has variations in the DNA-binding affinity among its binding motifs, with SBM being stronger than the non-bipartite motifs, including RGTGCA.

We further explored SBMs in the DIP peak regions to evaluate whether SBMs are recognized by the tandem AP2 domains of PbSIP2. The analysis detected 129 and 737 peaks containing fSBM and mSBM (25% of the DIP-seq peaks in total), respectively (Fig 6G). Furthermore, average fold enrichment values of fSBM and mSBM peaks were higher than that of the other peaks (*p*-values = 4.3 × 10^−4^ and 1.8 × 10^−3^, respectively), similar to the ChIP-seq results (Fig 6H). These results suggest that the variable DNA-binding affinities observed for PbSIP2 ChIP-seq were determined by its tandem AP2 domains.

Among the PbSIP2 target genes, 175 genes had a SBM peak on their upstream. These SBM-associated targets included 15 rhoptry genes and a few other merozoite-related genes, such as *gap45* and *msp10* (Fig 6I). Rhoptry genes were also detected in genes that were highly downregulated in *pbsip2*-cKO^Rapa+^ (Fig 4D), suggesting a relationship between genes associated with SBM and high transcription levels induced by PbSIP2. Consistently, among the downregulated genes, those containing the SBM within 1200 bp from their start codon tended to have higher −log_2_(fold change) values compared with the other downregulated genes (Fig 6J). Collectively, these results indicate that PbSIP2 acts as a stronger *trans*-activating factor on the SBM than on the other motifs, as it preferentially binds to the SBM.

## Discussion

Schizogony is a crucial step in the intraerythrocytic proliferation of *Plasmodium*. This process proceeds in two major steps: first, the parasites undergo several asynchronous rounds of nuclear division to form a multinucleated cell; second, multiple progeny merozoites are produced through segmentation (cytokinesis) simultaneously with a final round of karyokinesis. Merozoite components are expressed during the first step, and buds for each progeny merozoite (including rhoptries and apical ends) are already associated with the nuclei by the beginning of segmentation. Our results showed that PbSIP2 was expressed during the first step, and conditional knockout of *pbsip2* resulted in developmental arrest before the second step of schizogony. Furthermore, PbSIP2 broadly activated merozoite-related genes, including those involved in merozoite morphogenesis, motility, and invasion. These results suggest that PbSIP2 plays an essential role in schizogony as a master transcription factor that regulates merozoite formation. However, the role of PbSIP2 does not encompass the entire schizogony process. Notably, the *pbsip2*-knockout parasites were still capable of producing multinucleated cells, suggesting that PbSIP2 does not play a major role in DNA replication or nuclear division. Consistently, genes related to these processes were barely included among the PbSIP2 targets. Hence, other transcription factors should regulate the expression of such genes, and further exploration of these factors is required to better understand transcriptional regulation throughout the schizont development.

The ortholog of PbSIP2 in *P. falciparum*, PfSIP2, has been previously identified as a protein that binds to the SPE2 [(T/G)GTGC(A/G)] motif, which exists in tandem on the subtelomeric *var* gene promoters^16^. Based on these results, the authors suggested that PfSIP2 may be involved in chromosome end biology, such as chromosomal replication and segregation. Meanwhile, the authors also mentioned the possibility of other roles for PfSIP2 because subtelomeric SPE2 arrays are only detected in the *P. falciparum* genome. Our results showed that PbSIP2 functions as a *trans*-regulatory factor for merozoite-related genes through its association with motifs similar to SPE2 (GTGCA and SBM). These motifs are analogous to putative *cis*-regulatory elements upstream of invasion-related genes previously identified in the *P. falciparum* genome^11,12^. Taken together, we believe that the primary role of SIP2 in *Plasmodium* is the activation of merozoite-related genes through association with these specific *cis*-regulatory elements and promotion of merozoite formation. In this study, we demonstrated that SIP2 is particularly conserved in parasites belonging to the orders Haemosporida and Piroplasmida. Alignment of their AA sequences showed considerably high similarity within the tandem AP2 domains, including the linker regions, suggesting that their roles may also be conserved. Among the apicomplexan parasites, only those in Haemosporida and Piroplasmida produce an invasive stage that can infect host RBCs. Thus, SIP2 may play a conserved role as a master regulator of RBC-invasive stage development in these parasites and may have an evolutionary origin from their common ancestor.

ChIP-seq and DIP-seq analyses revealed that PbSIP2 binds to GTGCA and has a stronger affinity for RGTGCA. Moreover, PbSIP2 was more strongly associated with the SBM. The 3D structure of the tandem AP2 domains of PbSIP2 showed parallel AP2 domains with a distance that almost matched the distance between the two motifs in the SBM on B-DNA. Thus, the DNA-binding configuration of PbSIP2 could be considered as follows: one AP2 domain binds to GTGCA, and its binding becomes stronger when the motif is RGTGCA. Furthermore, when the other AP2 domain additionally binds to 5’-side TGCA in the SBM, PbSIP2 can be most strongly associated with DNA. Campbell et al. previously reported that in the protein-binding microarrays for the AP2 domains of PfSIP2, the N-terminal-side AP2 domain alone was capable of binding to the GGTGCA motif^35^. Thus, the N-terminal AP2 may be responsible for binding to GTGCA in the SBM. We propose that due to these different DNA-binding properties of the tandem AP2 domains, variations in transcription levels can be established among PbSIP2 targets. The *Plasmodium* genome encodes approximately 30 sequence-specific transcription factor genes, which seems to be a small number for eukaryotic species with a complex life cycle. Considering this, the parasite might use the above mechanism to control variable transcription levels of stage-specific genes with a small number of *trans*-regulatory factors.

During *Plasmodium* IDC, gene transcription is periodically regulated to generate a simple cascade of stage-specific transcriptomes^7^. AP2 transcription factors are also periodically expressed during IDC^35^, and some are essential for the asexual blood-stage development. These suggest that the transcriptional activators responsible for stage-specific transcription during IDC could be identified in these AP2 transcription factors. Nevertheless, the involvement of AP2 transcription factors in stage-specific transcription during IDC has barely been elucidated, and most AP2 transcription factors investigated in previous studies have been reported to be involved in the silencing of subtelomeric multigenes^36–39^. As a rare example, PfAP2-I has been shown to be a transcriptional activator of invasion-related genes during the asexual blood-stage development^40^. However, the activation of invasion-related genes does not seem to be the major role of PfAP2-I during IDC because the expression of PfAP2-I peaks in the trophozoite stage and decreases during the schizont stage. Moreover, the target genes of PfAP2-I include only a limited number of invasion-related genes. We believe that further investigation is required to evaluate other roles of PfAP2-I, such as ChIP-seq analyses at its peak expression period and conditional knockout analyses to assess the impact of disrupting *ap2-i* in parasite asexual development.

Studies on transcriptional regulators in *Plasmodium* have demonstrated surprisingly simple mechanisms of stage-specific transcriptional regulation throughout the life cycle. Of note, these studies have advanced our understanding of the mechanisms regulating sexual development and formation of the two other invasive stages in *Plasmodium*, namely sporozoites and ookinetes^29–31,41–45^. Our data indicate that merozoite formation is also regulated by such simple transcriptional regulation through the function of PbSIP2. We believe that further investigation of other AP2 transcription factors, especially those essential for asexual blood-stage development, would enable us to understand the entire *Plasmodium* IDC at the transcriptional level; *i.e.* a cascade of stage-specific transcription factors and their target genes explain the periodic gene transcription during IDC.

## Materials and methods

### Ethical statement

All experiments in this study were performed according to the recommendations in the Guide for the Care and Use of Laboratory Animals of the National Institutes of Health to minimize animal suffering and were approved by the Animal Research Ethics Committee of Mie University (permit number 23–29).

### Parasite preparation

All the parasites used in this study were inoculated into ddY mice. The transgenic parasites generated in this study were derived from *P. berghei* ANKA strain. Schizont culture was performed at 37 °C using RPMI1640 medium supplemented with 25% fetal bovine serum (Cosmo Bio), 100 units/mL penicillin, and 100 μg/mL streptomycin (Gibco). For cell cycle synchronization, mature schizonts produced by culturing for 16 h were harvested by density gradient centrifugation using an iodixanol solution (Optiprep, Serumwerk Bernburg) mixed with tricine solution to adjust the density to 1.077 g/mL. The harvested schizonts were intravenously injected into mice.

### Generation of mutant parasites

For tagging PbSIP2 with GFP, the DNA sequences of the two homologous regions were cloned into the *gfp*-fusion vector, which has an *hdhfr* expression cassette next to *gfp* as a pyrimethamine-selectable marker^46^, to fuse *pbsip2* in frame with *gfp*. The plasmid was linearized by *Xho*I and *Not*I digestion before use in transfection experiments. For gene editing with the CRISPR/Cas9 system using PbCas9, donor DNAs were constructed by overlap PCR, cloned into pBluescript KS (+) (Addgene) using the *Xho*I and *Bam*HI sites by In-Fusion cloning, and then amplified by PCR from the constructed plasmid. The sgRNA target sites were designed using CHOPCHOP (https://chopchop.cbu.uib.no/), and sgRNA vectors were constructed as previously described^23^.

Transfection was performed using the Amaxa Basic Parasite Nucleofector Kit 2 (LONZA). All transfectants were selected by treatment of mice with 70 μg/mL pyrimethamine in their drinking water. Recombination was confirmed by PCR and/or Sanger sequencing, and clonal parasites were obtained by limiting dilution. All primers used in this study are listed in Table S4.

### Fluorescence analysis

Fluorescence analysis was performed using Olympus BX51 microscope with Olympus DP74 camera. Nuclei were stained by incubating infected blood in PBS with 1 ng/mL Hoechst 33342 for 10 min at 37 °C before analysis.

### Conditional knockout of *pbsip2*-cKO parasites

*pbsip2*-cKO parasites were cultured for 16 h, and mature schizonts were harvested by density gradient centrifugation and inoculated into mice. Six hours after inoculation, the whole blood was harvested and split into two cultures. At the beginning of the culture, 1/20,000 volume of rapamycin (Wako) solution (200 μM DMSO stock) was added to one culture, achieving the final concentration of 10 nM, (*pbsip2*-cKO^Rapa+^) and the same volume of DMSO was added to the other (*pbsip2*-cKO^Rapa−^). For genotyping PCR, genomic DNAs were harvested from 100 μL of each culture. The primers used are listed in Table S4.

### ChIP-seq and sequencing data analysis

The cell cycle of PbSIP2::GFP parasites was synchronized as described above and cultured from 6 h post-injection (hpi). At 24 hpi, the cultures were passed through a Plasmodipur filter and were immediately fixed in 1% formalin. After fixing for 1 h at 30 °C, RBCs were lysed in ice-cold 150 mM NH_4_Cl solution, and then the residual cells were lysed in SDS lysis buffer (50 mM Tris-HCl, 1% SDS, 10 mM EDTA). The cell lysate was sonicated using the Bioruptor (Cosmo Bio) to shear the chromatin and mixed with anti-GFP polyclonal antibodies (ab290, Abcam) conjugated to Protein A Magnetic Beads (Invitrogen). DNA fragments were purified from the immunoprecipitated chromatin and subjected to library construction using the KAPA HyperPrep Kit (Kapa Biosystems). The library was sequenced using the Illumina NextSeq 500. Before immunoprecipitation, DNA fragments were purified from an aliquot of the cell lysate and sequenced to obtain the input sequence data. Two biologically independent experiments were performed for the data analysis.

Sequence data were mapped onto the reference genome sequence of *P. berghei* (v3.0, downloaded from PlasmoDB 46) using Bowtie 2. Reads aligned onto more than two sites were removed from the mapping data, and peaks were called by macs2 callpeak with fold enrichment > 2.5 and *q*-value < 0.01 using input sequence data as a control. The parameters for all programs were set to default, unless otherwise indicated. Common peaks in duplicates were defined as those with a distance of less than 150 bp between the peak summits. The enrichment of motifs within 50 bp of the peak summits was analyzed using Fisher’s exact test. Genes with peaks 1200 bp upstream from their start codons were identified as target genes.

### RNA-seq and sequence data analysis

*pbsip2*-cKO^Rapa−^ and *pbsip2*-cKO^Rapa+^ were prepared as described in the conditional knockout method above. The cultures were passed through a Plasmodipur filter at 26 hpi, and RBCs were lysed in ice-cold 150 mM NH_4_Cl solution. Total RNA was extracted from the parasites using Isogen II reagent (Nippon Gene), and RNA-seq libraries were prepared using the KAPA mRNA HyperPrep Kit (Kapa Biosystems) for each sample. Libraries were sequenced using the MGI DNBSEQ-G400 (MGI Tech Co., Ltd.). Three biologically independent experiments were performed for each sample. The sequence data were mapped onto the reference *P. berghei* ANKA v3 genome using HISAT2, with the maximum intron length of 1000. The number of reads mapped onto each gene was calculated using featureCounts and compared using DESeq2. Genes in the subtelomeric regions were excluded from the differential expression analysis. The parameters for all programs were set to default, unless otherwise indicated.

### Preparation of recombinant protein and DIP-seq analysis

Recombinant proteins were prepared as previously described^44^. Briefly, the DNA sequence encoding the tandem AP2 domains of PbSIP2 (position 87–214) was cloned into the MBP-fusion vector, pMal-c5X (NEB), using the *Not*I and *Bam*HI sites, and the plasmid was introduced into *E. coli* strain DH5α. The transformed *E. coli* was cultured for 12 h at 37 °C, and then expression of the MBP-fused protein was induced by isopropyl β-D-thiogalactopyranoside (final concentration of 200 nM) in the culture. Recombinant protein of AP2 domains fused with MBP (MBP::PbSIP2) was purified using amylose resin (NEB) and recovered in 10 mM maltose solution.

The MBP::PbSIP2 (5 μg) was mixed with *P. berghei* ANKA genomic DNA fragments (2 μg) in 400 μL of Binding Buffer (100 mM KCl, 2m M MgCl_2_, 2 mM Tris-HCl, 10 μM ZnSO_4_, and 10% glycerol) and incubated for 30 min. The recombinant proteins and bound DNA fragments were purified using amylose resin. DNA fragments were subjected to library preparation and NGS using the Illumina NextSeq 500, as described for the ChIP-seq analysis. Genomic DNA fragments before use for DIP were sequenced as the input. Analysis of the sequence data was performed as described for ChIP-seq, except for the peak calling parameters (fold enrichment > 3.0, *q*-value < 0.01).

### ChIP-qPCR and RT-qPCR for reporter experiments

ChIP-qPCR and RT-qPCR were performed using PbSIP2::GFP^Cas9^ and PbSIP2::GFP^Cas9_cismut^ to assess the function of the PbSIP2 binding motif as a *cis*-regulatory motif. For ChIP-qPCR analysis, ChIP experiments were performed at 24 hpi, as described for the ChIP-seq analysis. ChIP and input samples were each prepared using 200 and 10 μL of sonicated cell lysate, respectively. For RT-qPCR analyses, total RNA was harvested at 26 hpi using the Isogen II reagent (Nippon Gene), and cDNA was synthesized from the total RNA using the PrimeScript RT reagent Kit with gDNA Eraser (Takara). Quantification of ChIPed DNA and cDNA was performed by real-time qPCR using the TB Green Fast qPCR Mix (Takara) and Thermal Cycler Dice Real Time System II (Takara). Amplification was performed for 40 cycles, and the cycle threshold (Ct) was detected between 20 and 35 cycles. For ChIP-qPCR analysis, %input values were calculated as 2^(Ct^ChIP^-Ct^input^) × (1/20) × 100. Three biologically independent samples were prepared for each experiment and used for analysis. All primers used are listed in Table S4.

## Data availability

All FASTQ files for ChIP-sequencing, RNA-sequencing and DIP-sequencing experiments are available from the Gene Expression Omnibus database (accession numbers GSE266502, GSE266503, and GSE266869).

## Acknowledgements

The authors are grateful to Asami Noro (Mie University) for technical assistance.

## Supporting information

**S1 Fig.**
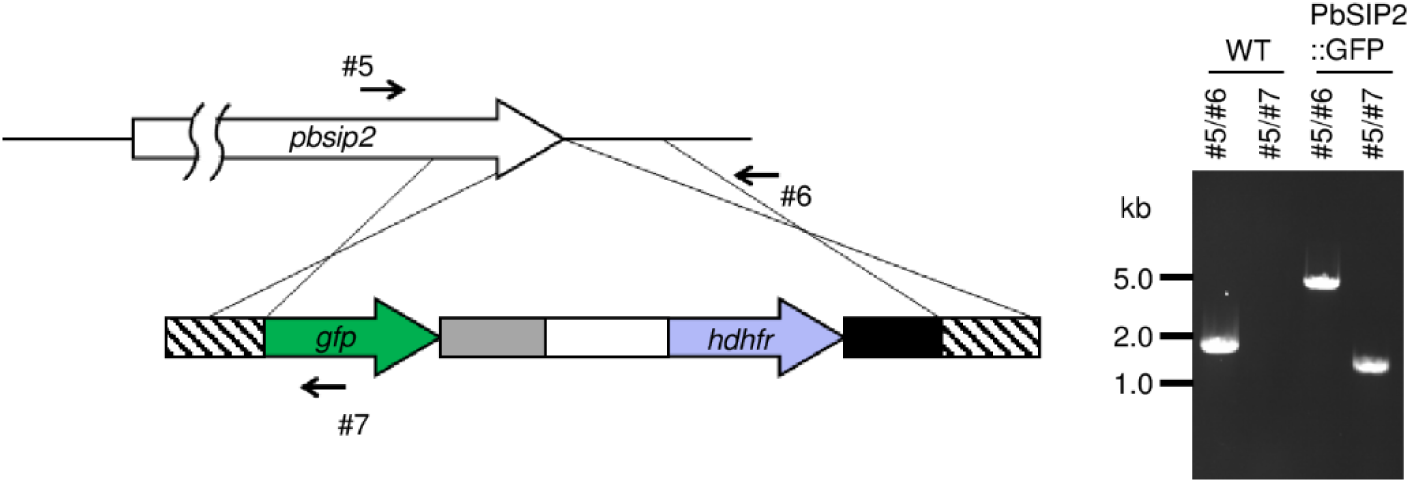
Genotyping of PbSIP2::GFP. Schematic illustration of gene editing at the *pbsip2* locus is shown on the left. A gel image from the genotyping PCR analysis is shown on the right side. The primer numbers are listed in S4 Table.

**S2 Fig.**
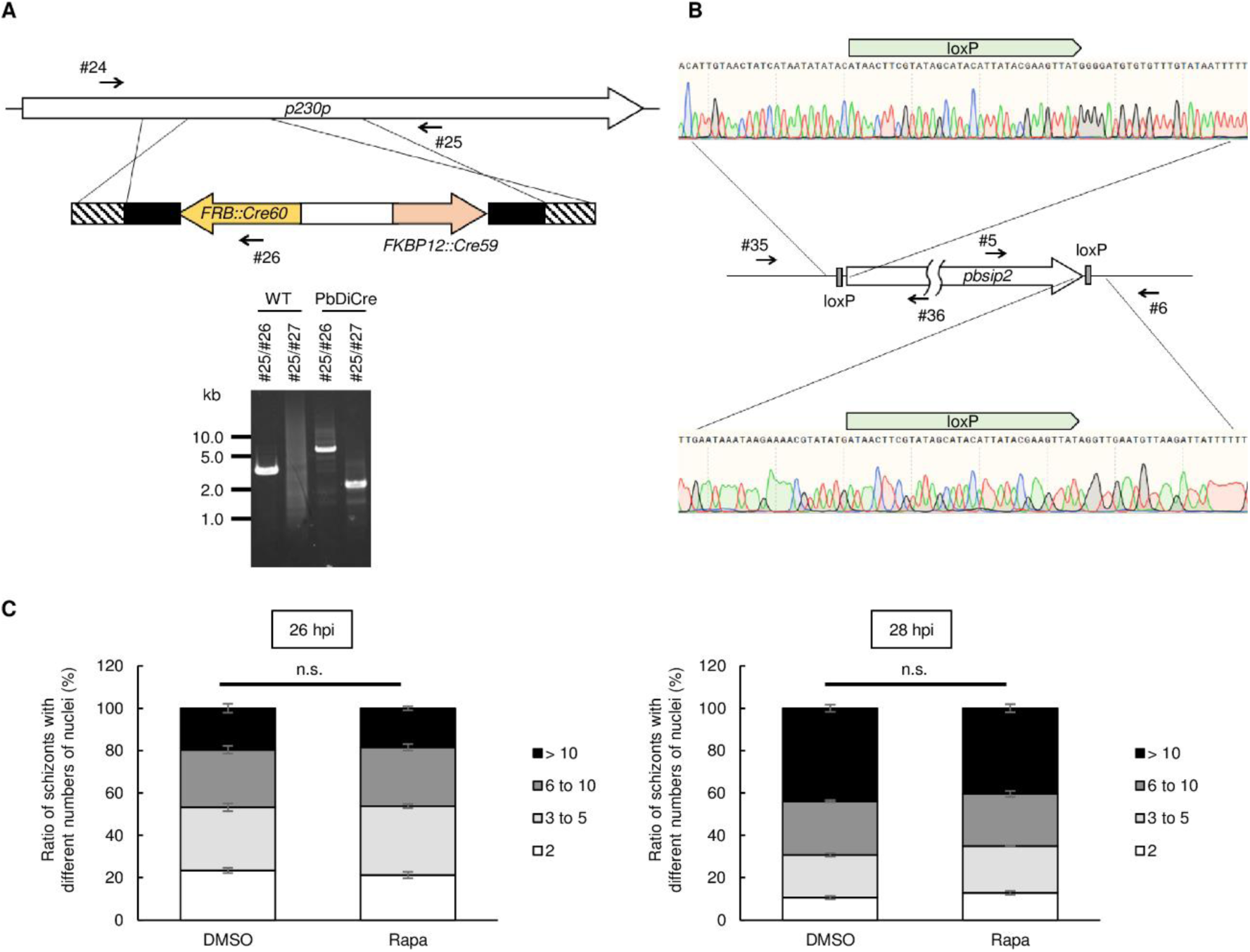
DiCre-mediated conditional knockout of *pbsip2*. (A) Genotyping of PbDiCre. Schematic illustration of gene editing at the *p230p* locus is shown at the top. A gel image of genotyping PCR analysis is shown at the bottom. The primer numbers are listed in S4 Table. (B) Genotyping of *pbsip2*-cKO. A schematic illustration of the *pbsip2* locus in *pbsip2*-cKO is shown. Sangar sequence results confirming insertion of loxP at 5’- and 3’-side of *pbsip2* are shown on the top and bottom, respectively. (C) Ratio of schizonts with different numbers of nuclei for PbDiCre in the absence and presence of rapamycin at 26 hpi (left) and 28 hpi (right). Error bars indicate the standard error of the mean values from three independent experiments. The *p*-values were calculated using a two-tailed Student’s t-test.

**S3 Fig.**
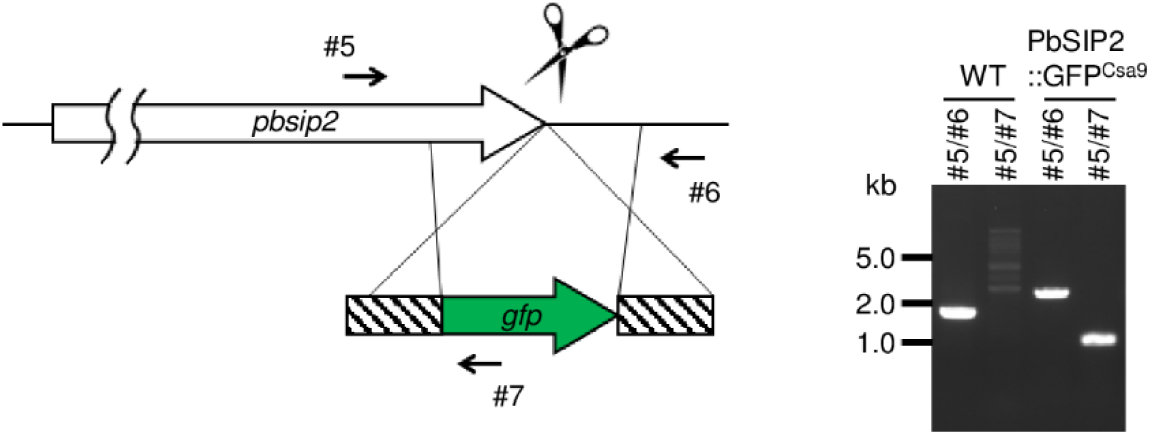
Genotyping of PbSIP2::GFP^Cas9^. Schematic illustration of gene editing at the *pbsip2* locus is shown on the left. A gel image from the genotyping PCR analysis is shown on the right side. The primer numbers are listed in S4 Table.

**S1 Table. ChIP-seq analysis of PbSIP2.** (A) Peaks identified in experiment 1. (B) Peaks identified in experiment 2. (C) Target genes of PbSIP2.

**S2 Table. Differential expression analysis between *pbsip2*-cKORapa− and *pbsip2*- cKORapa+.**

**S3 Table. DIP-seq analysis using MBP::PbSIP2.**

**S4 Table. List of primers used in this study.**

